# Conditioning a collective avoidance response in rummy-nose tetra

**DOI:** 10.1101/858704

**Authors:** Valentin Lecheval, Charlotte K. Hemelrijk, Guy Theraulaz

## Abstract

We develop an experimental method to induce controlled and local perturbations in a group of fish. Using the paradigm of the shuttle box we condition aversive escape reactions in groups of rummy-nose tetra (*Hemigrammus rhodostomus*) in response to a green light. Our experiments suggest that aversive conditioning can (i) be used successfully in this schooling species, (ii) trigger collective escape reactions and (iii) be transferred from the training set-up to a new environment. These results are discussed in the context of propagation of information among group members in reaction to external stimuli such as perturbations mimicking sudden changes in the environment, e.g. predator attack.

## 1. Introduction

Groups of rummy-nose tetra (*Hemigrammus sp.*) are highly coordinated (Faucher et al., 2010; Lecheval et al., 2018). Such groups of fish, in which individuals have aggregative tendencies and adopt the same orientation, are referred to as *schools* (Delcourt and Poncin, 2012). Fish schools are of particular interest for investigating collective behaviour of groups of animals in various ecological contexts and analysing how they adapt to challenging situations, such as predator attacks. When a school reacts collectively to an external perturbation, individuals react either directly to the perturbation itself or to the response of their neighbours (Domenici and Batty, 1994). Therefore, the source of information individuals respond to may differ among group members. Several techniques using virtual reality or a robotic dummy have been used to examine the impact of a single or few controlled virtual or robotic fish on collective behaviour (Faria et al., 2010; Polverino et al., 2012; Butail et al., 2013; Jouary et al., 2016; Bonnet et al., 2016; Stowers et al., 2017; Bonnet et al., 2018). Here we used a procedure of aversive conditioning to investigate how different reactions of individuals propagate in a group and affect the collective behaviour of a school of fish. We controlled the behaviour of some individuals in a group of *Hemigrammus rhodostomus* following a method used previously to investigate the propagation of information in sheep herds (Pillot et al., 2010, 2011; Miller et al., 2013; Toulet et al., 2015) or fish (Strandburg-Peshkin et al., 2013; Ioannou et al., 2015). However, in contrast with these studies focusing on appetitive conditioning to investigate propagation of information in the context of foraging, we use aversive conditioning by which an initially neutral stimulus becomes aversive after being repeatedly paired with an unconditioned aversive stimulus. Small electric shocks are associated with the emission of a particular light following the classical paradigm of the shuttle box, already tested in other species of fish (Horner et al., 1961; Woodard and Bitterman, 1973; Piront and Schmidt, 1988; Portavella et al., 2004; Pradel et al., 1999; Xu et al., 2007; Agetsuma et al., 2012). The aversive conditioning consists in training fish to perform fast-start escape responses when a green light is turned on. Inducing and controlling such escape responses in moving animal groups is of particular interest to better understand the propagation of information in the group when a single or a few individuals spot a predator in their neighbourhood. Previous studies with aversive conditioning in fish have been conducted in species that do not exhibit schooling behaviour, e.g. in zebrafish (*Danio rerio*) (Agetsuma et al., 2012) or goldfish (*Carassius auratus*) (Portavella et al., 2004). In the rummy-nose tetra, we show (i) that individuals learn to exhibit an escape response upon switching on a green light and (ii) that this conditioned escape response can be transferred to a new experimental environment.

## 2. Material and methods

Our fish experiments have been approved by the Ethics Committee for Animal Experimentation of the Toulouse Research Federation in Biology N ° 1 and comply with the European legislation for animal welfare.

### 2.1. Animals

A group of 70 rummy-nose tetras (*Hemigrammus rhodostomus*) were purchased from Amazonie Labège (http://www.amazonie.com) in Toulouse, France. The rummy-nose tetra is a tropical freshwater species. Fish were kept in 150 L aquariums on a 12:12 hour, dark:light photoperiod, at 27.5° C (±0.8° C) and were fed *ad libitum* with fish flakes.

### 2.2. Experiment 1: avoidance conditioning in a shuttle box

#### 2.2.1. Conditioned and naïve fish

Experiments with zebrafish have shown that when in group, fish may learn faster and be more active (Gleason et al., 1977). We thus conducted the conditioning experiments with a group of 6 fish – thereafter called *conditioned fish*. The fish have been randomly sampled and kept in a special breeding tank, under the same conditions as the other fish. The non conditioned individuals are labelled as *naïve fish*.

#### 2.2.2. Conditioning apparatus

Fish were trained using a shuttle box modified after Horner et al. (1961). The shuttle box (46 × 23 × 21 cm) is made of white plastic (polyvinyl chloride, PVC) (Figure 1A). In two compartments (20 × 15 × 20 cm) there are 84 green light-emitting diodes (LEDs) on 4 rows of 7 LEDs at each of the three walls in each compartment. LEDs of one compartment can be turned on independently from those of the other compartment (Figure 1A, inset). Compartments are separated by a trapezoidal hurdle 7.5 cm high. Two metallic plates conduct electricity in each compartment (7 V, 2.7 mA, measured on the electrodes, set to be the minimum electric power required to elicit a visible reaction from the fish). Following policies of animal welfare regarding environmental enrichment, we have filled the compartment with white gravel and white ceramic balls to a height of 4.5 cm. The water comprised 50% of water purified by reverse osmosis and 50% of water treated by activated carbon, heated at 27° C. The water level is set at 3 cm above the hurdle (i.e. 10.5 cm in each compartment). Light and electric stimuli are both delivered manually. Training sessions are monitored by a webcam (Logitech QuickCam Pro 9000).

**Figure 1:**
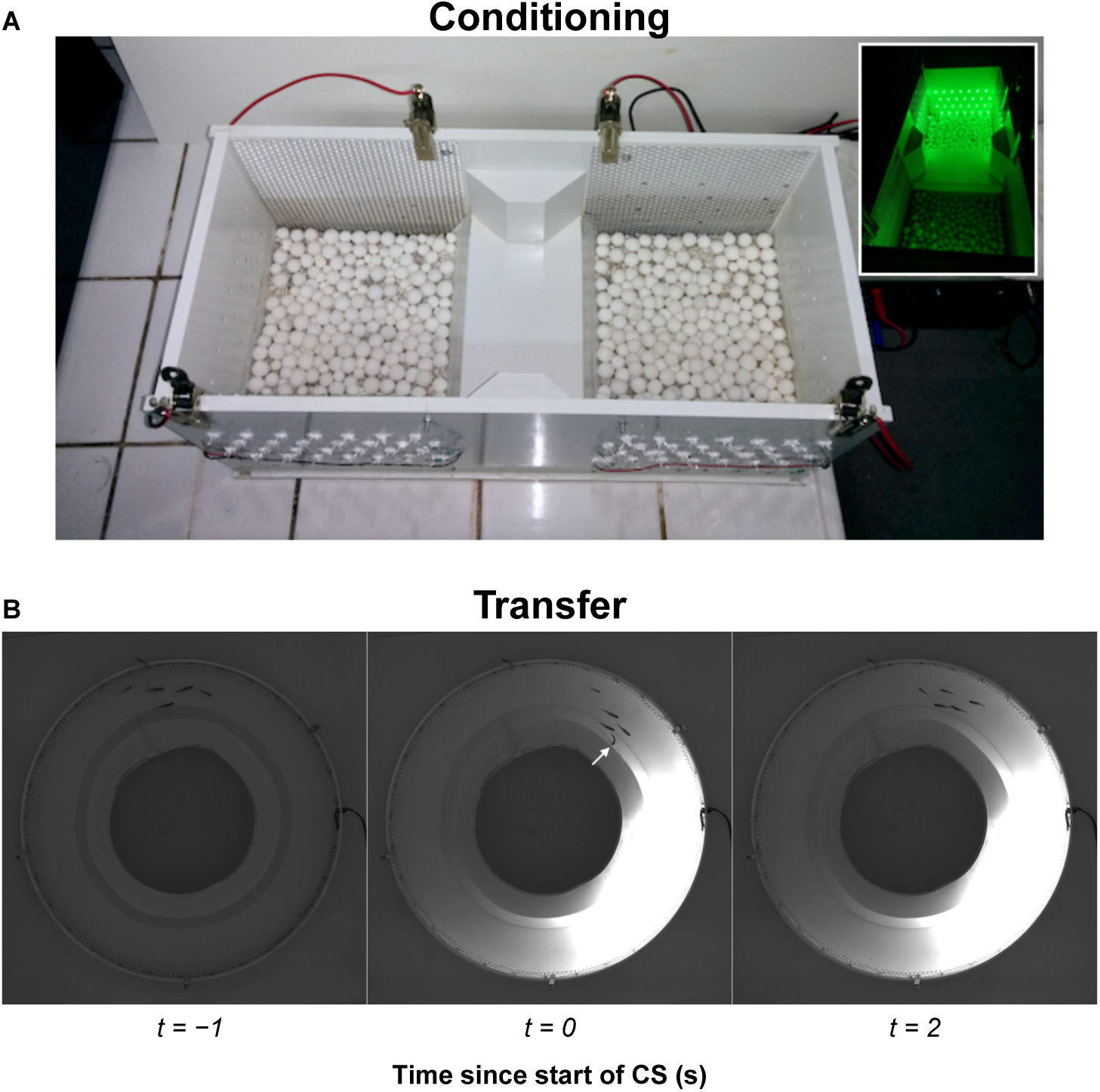
A). Pictures of the shuttle box (46 × 23 × 21 cm) used for the aversive conditioning (Experiment 1), with CS off and on (inset) in one compartment. B). Snapshots from an escape behaviour of a group with 5 conditioned fish during Experiment 2. Snapshots are taken from 1 second before the onset of the CS to 2 seconds after. The white arrow on the middle panel indicates the fish that responds first to the CS.

#### 2.2.3. Conditioning procedure

The group of 6 conditioned fish is placed in the setup for 15 min for familiarization and remains there during 5 min after the experiment is completed. Water is refreshed after each session.

#### 2.2.4. Acquisition

Training sessions were performed on a regular basis (77 sessions in total) with the same 6 fish^1^. Each training session consisted of 20 trials (intertrial time of 2 min). In each trial, at least one of the 6 fish had to cross the hurdle to avoid mild electric shocks (unconditioned stimulus, US) administered via electrodes 3 s^2^ after onset of a light signal (green light, conditioned stimulus, CS). The light signal was always turned on in the compartment where the majority of the individuals were located. A trial is labelled as a *Success* if at least one fish crossed the hurdle before the onset of electric shocks (i.e. within 3 s after onset of the CS) and *Failed* if not. If there is no fish crossing the hurdle within 15 s after the onset of electric shocks, all stimuli (CS + US) are turned off and the trial is labelled as *Failed*. The proportion of trials labelled as *Success* over the 20 trials will be referred to as the *relative frequency of escapes*.

Ten training sessions without US have been conducted on groups with 6 naïve fish randomly sampled from the breeding tank to measure the proportion of escapes due to the spontaneous exploration of the apparatus or due to the effect of green light prior to conditioning (Control).

Our dataset consists in the output (*Failed* or *Success*) of 200 trials for naïve fish and 1540 trials for conditioned fish.

### 2.3. Experiment 2: test in a new environment

#### 2.3.1. Experimental set-up

We used a rectangular experimental tank of size 120 × 120 cm, made of glass, supported by a structure of metal beam 20 cm high (Lecheval et al., 2018). A plywood plate was interposed between the mesh and the basin to dampen the forces exerted on the glass basin by its own weight and water. This structure also enables the attenuation of vibrations. The setup was placed in a chamber made by four opaque white curtains surrounded by four LED light panels to provide an isotropic lighting. Inside the experimental tank, a ring-shaped corridor was filled with 7 cm of water of controlled quality (50% of water purified by reverse osmosis and 50% of water treated by activated carbon) heated at 28.8° C (±0.7° C). The corridor was 10 cm wide with a circular outer wall of radius 35 cm. The shape of the circular inner wall was conic and its radius at the bottom was 25 cm. The conic shape was chosen to avoid the occlusion on videos of fish swimming too close to the inner wall. The outer wall has 2 rows of the same LEDs as used in the shuttle box, equally spaced by 1 cm. Only 35 cm of the entire diameter (thus 70 LEDs) can be turned on (CS).

#### 2.3.2. Experimental procedure

The experimental tests were conducted after the conditioning experiments. Fish were trained for 10 trials in the conditioning experiment a few hours before an experiment was done in the new set-up. Each fish was only used in a single experiment per day. Experiments concerned groups of 5 fish and were performed in the ring-shaped tank. Trajectories were recorded by a high-speed camera (R&D Vision) from above the set-up recording at 45 Hz (intertrial) or 180 Hz (trial), in high resolution (2000 × 2048 pixels). Groups are let to habituate for 20 min, with trajectories of individuals recorded at 45 Hz. At the onset of the CS, the camera automatically switches to record fish positions at 180 Hz. The CS is turned on for 3 s and fish trajectories are recorded subsequently for 30 s, still at 180 Hz. Several trials are performed on each group of fish, with an intertrial duration of 1 min in which the camera switches back at 45 Hz, before the next CS. Two conditions are tested, the *Control* condition, with 5 naïve fish (11 replicates, 5 trials per replicate) and the *Test* condition with 5 conditioned fish (2 replicates, 10 trials per replicate).

For each trial, we measure the number of fish that turn at least once in the direction opposite to the CS in the first 3 s following the onset of the CS (i.e while the light is on) and during the 3 s seconds after the light turns off (i.e. we monitor fish behaviour for 6 seconds after the onset of the CS). These two quantities estimate the strength of the collective reaction: if all individuals react fast when the CS is delivered, we expect them to turn while the light is on. We also measure whether all individuals are swimming in the direction opposite to the CS 6 s after its onset or not. When this condition is met, we count one collective escape for the respective trial. This quantity is used to discriminate between non-coordinated behaviours (e.g. where all individuals make U-turns several times and eventually the group does not swim away from the CS) and collective escape responses (i.e all group members make one clear U-turn and the group swims away from the CS).

We removed trials in which not all fish were aligned in the same direction just before the CS is turned on (4 trials out of 55 in the *Control* condition and 0 out of 20 in the *Test* condition).

## 3. Results

### 3.1. Experiment 1

We use logistic regressions with R (R Core Team, 2016) to investigate the effect of the conditioning experiments on the escape response of fish. We model the probability *p* that the binary result *r* of each trial (*Failed* (*r* = 0) or *Success* (*r* = 1)) is 1 as a function of the index of the trial *i*, the number of days *d* since the beginning of the conditioning for the considered group and of *c* that indicates whether the group is a control (*c* = 0) or a conditioned one (*c* = 1). We evaluate the full model:

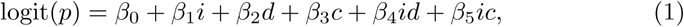

where *β*_*j*_, *j* ∈ [0, 6] are the regression coefficients. We search the model that minimises the Akaike information criterion (AIC) from the full model (AIC_full_ = 2231.43). The AIC addressed the trade-off between the goodness of fit of the model and the number *j* − 1 of explanatory variables. We find that it is not possible to discriminate between models with at least the three variable and one or two interaction terms (AIC for model without interaction is AIC_ni_ = 2233.23).

We report the exponential of the estimated regression coefficients (odd ratios) of the model without interaction

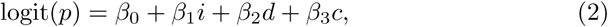

in Table 1. We note that models with interaction terms does not change our conclusions. We perform likelihood ratio tests for each of the three variables against a null model logit(*p*) = *β*_0_ to assess their significance in predicting the success or failure of a trial.

**Table 1:** Exponential of the estimated logistic regression coefficients (odd ratios and confidence intervals) of the model shown in Equation 2. We also report the *p*-values of the Wald statistic that tests for each *β*_*j*_ the null hypothesis *β*_*j*_ = 0 (no significant effect of the *j*^th^ variable). For *β*_1_ to *β*_3_, values greater than 1 indicate positive correlation while values less than 1 indicate negative correlation between *p* and the respective explanatory variable.

| | Estimate | 2.5% | 97.5% | $p$-value |
| --- | --- | --- | --- | --- |
| $\exp(\hat{\beta}_0)$ | 0.088 | 0.031 | 0.214 | |
| $\exp(\hat{\beta}_1)$ | 1.040 | 1.023 | 1.059 | 0.0054 |
| $\exp(\hat{\beta}_2)$ | 0.999 | 0.996 | 0.9995 | < 0.001 |
| $\exp(\hat{\beta}_3)$ | 11.683 | 7.548 | 18.782 | < 0.001 |

Our results show that *H. rhodostomus* can learn to escape quickly when a green light is turned on, in a social context. The proportion of escape responses (avoidance of the CS) is statistically larger in the conditioned group (0.57±0.02, mean±standard error, GLM: *D* = 155.27, *p*-value < 0.001) than in the control groups (0.125±0.03) (Figure 2A and 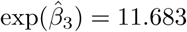, Table 1). Control groups reach a low proportion of escapes because they randomly swim in the shuttle box and may cross the barrier during the CS, either by accident or in reaction to the CS. We qualitatively assess an effect of the CS on the behaviour of the naïve fish which is different from the behaviour of conditioned fish. In general, groups of naïve fish become excited during CS but without any avoidance behaviour – they are even sometimes attracted towards the LEDs that emit the light. Within a training session, fish perform significantly better after several trials (GLM: *D* = 18.75, *p*-value < 0.001 and 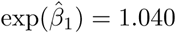). The time series of the proportion of successful responses of the conditioned group shows that the learning of the task occurs within a month (purple dots) and the best performances are achieved within the first five months (blue dots, Figure2A). From the seventh to the twelfth (green to yellow dots), despite 27 conditioning experiments, it was not possible to reach the performances obtained in the first months. This suggests that fish experience long-term habituation (GLM: *D* = 5.2051, *p*-value = 0.02 and 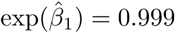), even though our protocol cannot properly test it (i.e. we should test the specificity of habituation to the stimulus to exclude an effect of fatigue due to the long period of time involved – a year, and the repeated use of electric shocks which might have affected fish). Although fish react less to the CS in the last sections of the time series, the achieved performance is, on average, still higher than the controls.

**Figure 2:**
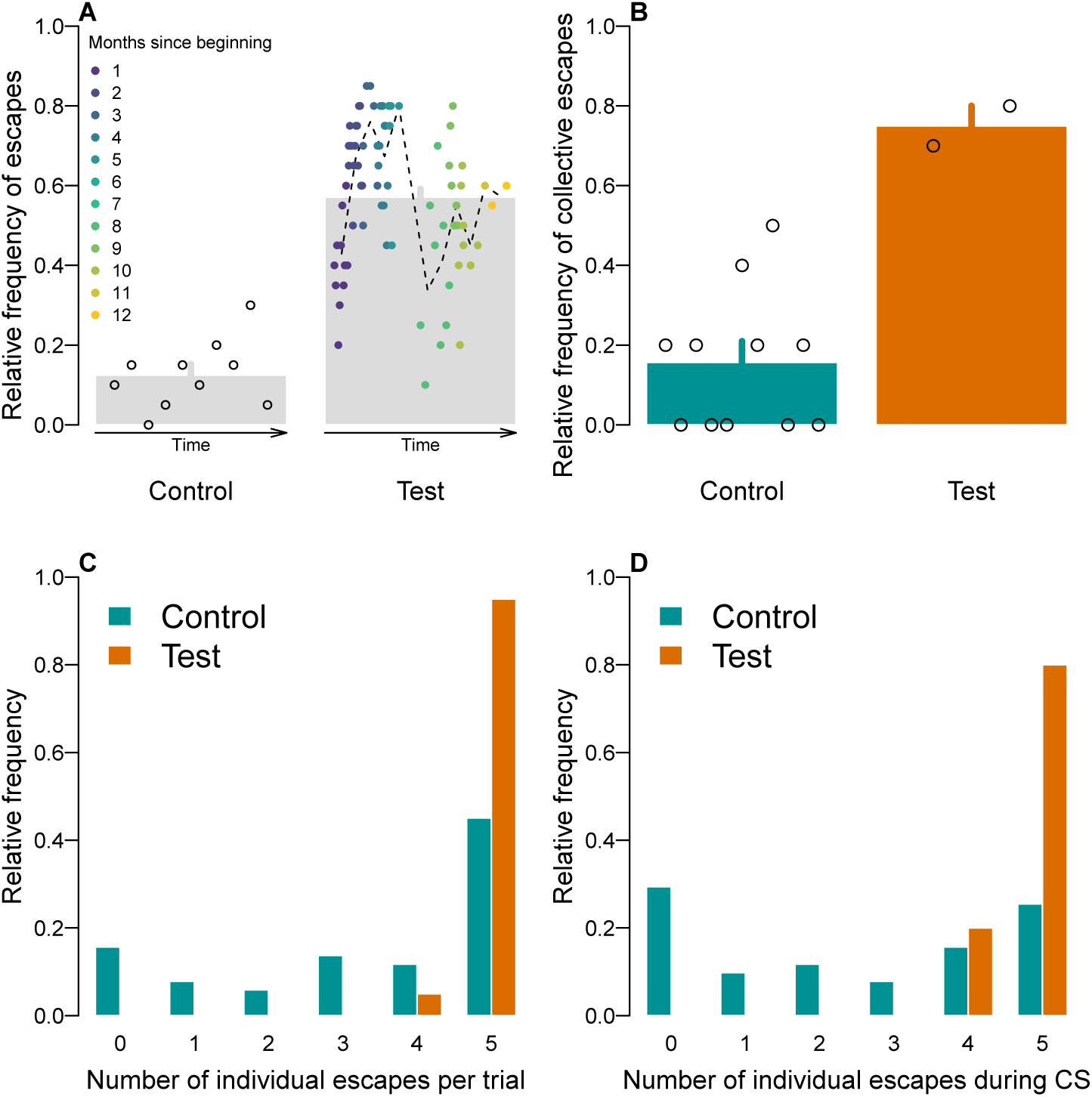
A). Learning performance in control and test conditions in Experiment 1. Open circles and dots show the proportion of success (at least one fish crossed the hurdle before the onset of electric shocks) of the group across all trials of each experiment as a function of time (from left (the first experiments) to right (the last experiments). The colour of the dots stand for the time since the beginning of the experiments in the *Test* condition. Dashed line is the monthly average, as a guide for the eye. Grey bars and grey lines stand respectively for mean and standard error across time. B-D). Results in Experiment 2 for each condition (*Control* for groups made of naïve fish and *Test* for groups of conditioned fish). B). Average relative frequency and standard error of collective escapes for each condition. We report results averaged over replicates (open circles). C). Distribution of individual escapes per trial, for each condition. D). Distribution of individual escapes per trial that occur during the CS (i.e. within 3 s), for each condition.

### 3.2. Experiment 2

Figure 1B shows a collective escape away from the CS performed by the group of conditioned fish: before the onset of the CS, the group is polarised (Figure 1B, *t* = −1 s); at the onset of the CS, one individual reacts instantaneously, performing a U-turn (*t* = 0 s) which propagates to the other group members and the group swims in the opposite direction in less than 2 s after the light is turned on.

We used R (R Core Team, 2016) and the packages `lme4` (Bates et al., 2015) and `lsmeans` (Lenth, 2016) to perform a binomial generalised linear mixed effects analysis of the occurrence of collective escapes in trials as a function of the condition of the replicate (*Control, Test*) and the index of the trial. Namely, we test (i) an effect of the condition (*Control* or *Test*) which could change the propensity to initiate a collective escape and (ii) an effect of habituation or fatigue to the CS across trials. As random effect, we use intercept for the experiment as well as by-experiment random slopes to account for the non-independence of the responses within a condition, since there are several trials per tested group of fish. *P* -values were obtained by likelihood ratio tests of the full model with the fixed effects against models with only one of each of the fixed effects. We also perform the likelihood ratio tests of the models with only one of each of the fixed effects against the null model with intercept and random effect only.

Both tests show a significant effect of the condition (*Control, Test*) on the occurrence of collective escapes and no significant effect of the trial (Table 2). We find that the CS is not neutral on the behaviour of the naïve groups. In 71% of all trials of *Control* condition, there are at least 3 fish that make individual U-turns (Figure 2C) but less than 50% turn within 3 s (Figure 2D). It results in naïve fish performing collective escapes in only 16% of all trials (Figure 2B). Instead, when all group members are conditioned (*Test* condition), groups perform collective escapes in 75% of all trials (Figure 2B). In that case, the response is in general collective and fast: in 80% of all trials all group members have turned within three seconds after the CS is turned on (Figure 2D). In short, although groups of naïve fish react to the CS by fluctuating headings of individuals, they react more slowly and not collectively, in contrast to the groups composed of conditioned fish.

**Table 2:** *P* -values of the likelihood ratio tests performed to assess the significance of the effects of the condition and of the trial to predict the occurrence of collective escapes from binomial generalised linear mixed effect models.

|  | Only Condition | Only Trial | Null Model |
| --- | --- | --- | --- |
| Full Model | 0.43 | 0.001** | 0.001** |
| Only Condition |  |  | < 0.001*** |
| Only Trial |  |  | 0.10 |

## 4. Discussion

Our experiments show that it is possible to train groups of *H. rhodostomus* to collectively escape from an unsafe zone in an aversive paradigm. We also show that this conditioned behaviour can be successively transferred to a new environment. The possibility to condition *H. rhodostomus* opens opportunities to investigate collective behaviour with obligate schoolers also, such as experiments in which the number of individuals informed of an appropriate response in reaction to a stimulus is controlled.

The duration of the CS before the onset of electric shocks (3 s) is very short compared to usage in previous studies (7.5 s (Agetsuma et al., 2012), 10 s (Horner et al., 1961; Woodard and Bitterman, 1973; Portavella et al., 2004), 12 s (Pradel et al., 1999; Xu et al., 2007), or 20 s (Piront and Schmidt, 1988, although with a different protocol)). This has to be taken into account to assess the conditioning success of *H. rhodostomus* (which was on average close to 63% during the first six months of conditioning) compared to the other studies, achieving learning criterion of 70% (Portavella et al., 2004) or 80% (Pradel et al., 1999; Piront and Schmidt, 1988). This choice of a duration of the CS before electric shocks shorter than in previous studies was made to elicit faster aversive reactions in response to the CS.

Our experimental results also suggest that the training leads to fatigue or habituation because the performance of the conditioned group decreased after the first 48 experiments conducted over a period of 4 months. Even if our protocol was not designed to test the presence of fatigue or habituation, our results suggest that this decrease in the performance of the conditioned groups was not a consequence of a lapse in fish memory: fish kept on performing fast aversive responses, although less often with time.

As for the set of experiments in the new environment, a ring-shaped tank and the propagation of the induced perturbation, we found that the occurrence of fast, collective escape increases when the group is made of conditioned fish. Given the time resolution of our videos (180 fps after the onset of the CS), data with the positions of fish will help to investigate qualitatively and quantitatively the collective behaviour and the propagation of information of groups reacting to a threat.

In the experiment, we aimed to control (i) which individual would react to a green light (by conditioning them), and (ii) how they would react to it (by escaping away from it). In the present case, by mixing naïve and conditioned fish, we did not entirely succeed to control the identity of the first responder because green light also has an effect on naïve fish. This needs to be fixed in future experiments, by either changing the conditioning stimulus (e.g. by reducing its intensity) or by familiarising the naïve fish to the green light. We did, however, manage to train fish to escape a green light collectively, thus controlling (to some extent) that the green light is perceived as a threat. These results motivate future use of this method to investigate how the proportion of informed individuals among group members affects the propagation of information in the whole group, in particular in the context of escapes.

## Footnotes

1 One of the 6 fish died, 10 months after the beginning of the experiment. The death did not occur during an experiment and no wounds were noticed. Experiments were thus performed with the other 5 fish thereafter.

2 The CS-US interval had first been set to 5 s and then to 3 s after the 7 first experiments, to elicit faster avoidance reactions.

## References

Agetsuma, M., Aoki, T., Aoki, R., Okamoto, H., 2012. Cued Fear Conditioning in Zebrafish (Danio rerio), in: Kalueff, A.V., Stewart, A.M. (Eds.), Zebrafish Protocols for Neurobehavioral Research. Humana Press, Totowa, NJ, pp. 257–264.

Bates, D., Mächler, M., Bolker, B., Walker, S., 2015. Fitting Linear Mixed-Effects Models Using lme4. Journal of Statistical Software 67, 1–48. doi:10.18637/jss.v067.i01.

Bonnet, F., Gribovskiy, A., Halloy, J., Mondada, F., 2018. Closed-loop interactions between a shoal of zebrafish and a group of robotic fish in a circular corridor. Swarm Intelligence 12, 227–244. URL: https://doi.org/10.1007/s11721-017-0153-6, doi:10.1007/s11721-017-0153-6.

Bonnet, F., Kato, Y., Halloy, J., Mondada, F., 2016. Infiltrating the zebrafish swarm: design, implementation and experimental tests of a miniature robotic fish lure for fish–robot interaction studies. Artificial Life and Robotics 21, 239–246. URL: https://doi.org/10.1007/s10015-016-0291-8, doi:10.1007/s10015-016-0291-8.

Butail, S., Bartolini, T., Porfiri, M., 2013. Collective Response of Zebrafish Shoals to a Free-Swimming Robotic Fish. PLOS ONE 8, e76123. URL: https://doi.org/10.1371/journal.pone.0076123, doi:10.1371/journal.pone.0076123.

Delcourt, J., Poncin, P., 2012. Shoals and schools: Back to the heuristic definitions and quantitative references. Reviews in Fish Biology and Fisheries 22, 595–619. doi:10.1007/s11160-012-9260-z.

Domenici, P., Batty, R.S., 1994. Escape manoeuvres of schooling Clupea harengus. Journal of Fish Biology 45, 97–110.

Faria, J.J., Dyer, J.R.G., Clément, R.O., Couzin, I.D., Holt, N., Ward, A.J.W., Waters, D., Krause, J., 2010. A novel method for investigating the collective behaviour of fish: introducing ‘Robofish’. Behavioral Ecology and Sociobiology 64, 1211–1218. URL: https://doi.org/10.1007/s00265-010-0988-y, doi:10.1007/s00265-010-0988-y.

Faucher, K., Parmentier, E., Becco, C., Vandewalle, N., Vandewalle, P., 2010. Fish lateral system is required for accurate control of shoaling behaviour. Animal Behaviour 79, 679–687. URL: http://www.sciencedirect.com/science/article/pii/S0003347209005788, doi:10.1016/j.anbehav.2009.12.020.

Gleason, P.E., Weber, P.G., Weber, S.P., 1977. Effect of group size on avoidance learning in zebra fish,Brachydanio rerio (Pisces: Cyprinidae). Animal Learning & Behavior 5, 213–216. doi:10.3758/BF03214081.

Horner, J.L., Longo, N., Bitterman, M.E., 1961. A Shuttle Box for Fish and a Control Circuit of General Applicability. The American Journal of Psychology 74, 114–120.

Ioannou, C.C., Singh, M., Couzin, I.D., 2015. Potential leaders trade off goal-oriented and socially oriented behavior in mobile animal groups. The American Naturalist 186, 284–293. URL: https://doi.org/10.1086/681988, doi:10.1086/681988, arXiv:https://doi.org/10.1086/681988. pMID: 26655156.

Jouary, A., Haudrechy, M., Candelier, R., Sumbre, G., 2016. A 2d virtual reality system for visual goal-driven navigation in zebrafish larvae. Scientific Reports 6, 34015. URL: https://doi.org/10.1038/srep34015.

Lecheval, V., Jiang, L., Tichit, P., Sire, C., Hemelrijk, C.K., Theraulaz, G., 2018. Social conformity and propagation of information in collective u-turns of fish schools. Proceedings of the Royal Society of London B: Biological Sciences 285. doi:10.1098/rspb.2018.0251.

Lenth, R.V., 2016. Least-Squares Means: The R Package lsmeans. Journal of Statistical Software 69, 1–33. doi:10.18637/jss.v069.i01.

Miller, N., Garnier, S., Hartnett, A.T., Couzin, I.D., 2013. Both information and social cohesion determine collective decisions in animal groups. Proceedings of the National Academy of Sciences 110, 5263–5268. doi:10.1073/pnas.1217513110.

Pillot, M.H., Gautrais, J., Arrufat, P., Couzin, I.D., Bon, R., Deneubourg, J.L., 2011. Scalable Rules for Coherent Group Motion in a Gregarious Vertebrate. PLOS ONE 6, 1–8. doi:10.1371/journal.pone.0014487.

Pillot, M.H., Gautrais, J., Gouello, J., Michelena, P., sibbald, A., Bon, R., 2010. Moving together: Incidental leaders and naïve followers. Behavioural Processes 83, 235 – 241. doi:http://dx.doi.org/10.1016/j.beproc.2009.11.006.

Piront, M.L., Schmidt, R., 1988. Inhibition of long-term memory formation by anti-ependymin antisera after active shock-avoidance learning in goldfish. Brain Research 442, 53 – 62. doi:https://doi.org/10.1016/0006-8993(88)91431-X.

Polverino, G., Abaid, N., Kopman, V., Macrì, S., Porfiri, M., 2012. Zebrafish response to robotic fish: preference experiments on isolated individuals and small shoals. Bioinspiration & Biomimetics 7, 036019. URL: http://dx.doi.org/10.1088/1748-3182/7/3/036019, doi:10.1088/1748-3182/7/3/036019.

Portavella, M., Torres, B., Salas, C., 2004. Avoidance Response in Goldfish: Emotional and Temporal Involvement of Medial and Lateral Telencephalic Pallium. Journal of Neuroscience 24, 2335–2342. doi:10.1523/JNEUROSCI.4930-03.2004.

Pradel, G., Schachner, M., Schmidt, R., 1999. Inhibition of memory consolidation by antibodies against cell adhesion molecules after active avoidance conditioning in zebrafish. Journal of Neurobiology 39, 197–206. doi:10.1002/(SICI)1097-4695(199905)39:2<197::AID-NEU4>3.0.CO;2-9.

R Core Team, 2016. R: A Language and Environment for Statistical Computing. R Foundation for Statistical Computing, Vienna, Austria.

Stowers, J.R., Hofbauer, M., Bastien, R., Griessner, J., Higgins, P., Farooqui, S., Fischer, R.M., Nowikovsky, K., Haubensak, W., Couzin, I.D., Tessmar-Raible, K., Straw, A.D., 2017. Virtual reality for freely moving animals. Nature Methods 14, 995–1002. URL: https://doi.org/10.1038/nmeth.4399, doi:10.1038/nmeth.4399.

Strandburg-Peshkin, A., Twomey, C.R., Bode, N.W., Kao, A.B., Katz, Y., Ioannou, C.C., Rosenthal, S.B., Torney, C.J., Wu, H.S., Levin, S.A., others, 2013. Visual sensory networks and effective information transfer in animal groups. Current Biology 23, R709–R711.

Toulet, S., Gautrais, J., Bon, R., Peruani, F., 2015. Imitation Combined with a Characteristic Stimulus Duration Results in Robust Collective Decision-Making. PLOS ONE 10, e0140188. doi:10.1371/journal.pone.0140188.

Woodard, W.T., Bitterman, M., 1973. Pavlovian analysis of avoidance conditioning in the goldfish (Carassius auratus). Journal of Comparative and physiological Psychology 82, 123.

Xu, X., Scott-Scheiern, T., Kempker, L., Simons, K., 2007. Active avoidance conditioning in zebrafish (Danio rerio). Neurobiology of Learning and Memory 87, 72 – 77. doi:https://doi.org/10.1016/j.nlm.2006.06.002.

